# Mouse B2 SINE elements function as IFN-inducible enhancers

**DOI:** 10.1101/2022.08.02.502523

**Authors:** Isabella Horton, Conor J. Kelly, David M. Simpson, Edward B. Chuong

## Abstract

Regulatory networks underlying innate immunity continually face selective pressures to adapt to new and evolving pathogens. Transposable elements (TEs) can affect immune gene expression as a source of inducible regulatory elements, but the significance of these elements in facilitating evolutionary diversification of innate immunity remains largely unexplored. Here, we investigated the mouse epigenomic response to type II interferon (IFN) signaling and discovered that elements from a subfamily of B2 SINE (B2_Mm2) contain STAT1 binding sites and function as IFN-inducible enhancers. CRISPR deletion experiments in mouse cells demonstrated that a B2_Mm2 element has been co-opted as an IFN-inducible enhancer of *Dicer1* and the nearby *Serpina3f* and *Serpina3g* genes. The rodent-specific B2 SINE family is highly abundant in the mouse genome and elements have been previously characterized to exhibit promoter, insulator, and non-coding RNA activity. Our work establishes a new role for B2 elements as inducible enhancer elements that influence mouse immunity and exemplifies how lineage-specific TEs can facilitate evolutionary divergence of innate immune regulatory networks.

## INTRODUCTION

The cellular innate immune response is the first line of defense against an infection, and is initiated by the activation of transcriptional networks that include antiviral and pro-inflammatory genes. While innate immune signaling pathways are generally conserved across mammalian species, the specific transcriptional networks are increasingly recognized to show differences across lineages (1, 2). These differences are widely attributed to independent evolutionary histories and continual selective pressures to adapt to new pathogens (3). Understanding how innate immune systems have evolved in different host genomes is critical for accurately characterizing and modeling responses that are related to autoimmunity or involved in disease susceptibility.

Transposable elements (TEs) are increasingly recognized as a source of genetic elements that shape the evolution of mammalian innate immune responses (1, 4). TE-derived sequences constitute roughly half of the genome content of most mammals, and are the predominant source of lineage-specific DNA. While most TE-derived sequences are degraded and presumed nonfunctional, TEs have occasionally been co-opted to function as genes or regulatory elements that benefit the host organism. In the context of host innate immunity, there are several reported examples of species-specific restriction factors that are encoded by TEs coopted for host defense, including Friend Virus 1, Syncytin, and Jaagsiekte sheep retrovirus (JSRV) (5–7). In many cases, TEs derived from ancient viral infections are poised for co-option since they already have the ability to bind to receptors, therefore blocking infection as a dominant negative mechanism (8).

More recently, TEs have also been identified as a source of non-coding regulatory elements that control inducible expression of cellular innate immunity genes (9). In the human genome, we previously showed that MER41 elements have been co-opted as enhancer elements to regulate multiple immune genes in human cells, including the *AIM2* inflammasome genes (10). Elements belonging to other transposon families, including LTR12, MER44, and THE1C, have also been co-opted to regulate inducible expression of immune genes (11–13). Notably, the majority of these families are primate-specific, supporting the co-option of TEs as a driver of primate-specific divergence of immune regulatory networks.

A key open question is whether the co-option of TEs as immune regulatory elements is evolutionarily widespread as a mechanism driving divergence of innate immune responses. Most research in this area has focused on human cells and primate-specific TE families, but different mammalian species harbor highly distinct and lineage-specific repertoires of TEs in their genomes. Due to the independent origin of most of these TEs in different species, it remains unclear whether the co-option of TEs is a rare or common mechanism contributing to the evolution of immune gene regulatory networks.

Here, we focused on the role of TEs in regulating murine innate immune responses. Mice are a commonly used model for human diseases but their immune system is appreciated to have significant differences. Transcriptomic studies have revealed that mouse and human immune transcriptomes show substantial divergence (2, 14), consistent with functional differences in inflammatory responses (15). The rodent and primate lineages diverged roughly 90 million years ago (16, 17) and 32% of the mouse genome consists of rodent-specific repeats (18). Therefore, we sought to define the potential role of TEs in shaping lineage-specific features of the murine innate immune response.

In our study, we re-analyzed transcriptomic and epigenomic datasets profiling the type II interferon (IFN) response in primary mouse macrophage cells. We screened for TEs showing epigenetic signatures of inducible regulatory activity, and identified a rodent-specific B2 subfamily as a substantial source of IFN-inducible regulatory elements in the mouse genome. As a case example, we used CRISPR to characterize a B2-derived IFN-inducible enhancer that regulates mouse *Dicer1.* These findings uncover a novel cis-regulatory role for the SINE B2 element in shaping the evolution of mouse-specific IFN responses.

## RESULTS

### Species-specific TEs shape the epigenomic response to type II IFN in mouse

To examine how TEs contribute to mouse type II IFN signaling regulation, we re-analyzed two independent transcriptomic and epigenomic datasets of primary bone marrow derived macrophages (BMDMs) that were stimulated with recombinant interferon gamma (IFNG) or untreated for 2 or 4 hours (19). These datasets included matched RNA-seq and ChIP-Seq for STAT1 and H3K27ac. The STAT1 transcription factor mediates the type II IFN response (20) by binding to enhancers and promoters containing the Gamma-IFN activation site (GAS) motif, and the H3K27ac modification is strongly associated with active enhancers (21). Using these datasets, we mapped both IFNG-inducible enhancers and IFNG-stimulated genes (ISGs).

Across both datasets, our analysis of the RNA-seq data identified a total of 1,896 ISGs (FDR < 0.05, log_2_ fold change > 1), which enriched for canonical genes associated with the IFNG response (GO:0034341, *p*-value = 9.394×10^-35^) (Supplemental Table S1). We predicted IFNG-inducible enhancers based on occupancy by the enhancer-associated histone mark H3K27ac and the transcription factor STAT1, which mediates type II IFN signaling. We identified 22921 regions bound by STAT1 in IFNG-induced cells, 18337 (80.0%) of which also resided within H3K27ac-enriched regions, indicating they are putative enhancers. Specificity of pulldown was confirmed by enrichment of canonical STAT1 binding motifs including the Gamma-IFN activation site (GAS) (E-value = 1.11×10^-746^) and IFN-stimulated response element (ISRE) (*p*-value = 1.01×10^-441^) motifs within the STAT1 ChIP-seq peaks (Supplemental Table S2).

Using this set of STAT1 binding sites, we next asked what fraction of binding sites were derived from mouse TEs. Using the summits of the STAT1 ChIP-seq peaks, we found that 26.6% resided within TEs, 71.1% of which contain significant matches (*p*-value < 1×10^-4^) to either ISRE or GAS motifs (Supplemental Table S3). These TEs likely represent direct binding sites of STAT1 with potential regulatory activity. We next asked whether any TE families were overrepresented within the set of predicted IFNG-inducible binding sites, using GIGGLE colocalization analysis (22). We identified three subfamilies enriched for STAT1 binding sites, including the rodent-specific B2_Mm2 subfamily (*p*-value = 7.18×10^-201^) as well as the RLTR30B_MM (*p*-value = 9.61×10^-77^) and RLTR30E_MM (*p*-value = 7.32×10^-31^) endogenous retrovirus subfamilies (Figure 1A, Supplemental Table S4). This indicates that the expansion of rodent-specific TE families has shaped the innate immune regulatory landscape in mouse.

**Figure 1.**
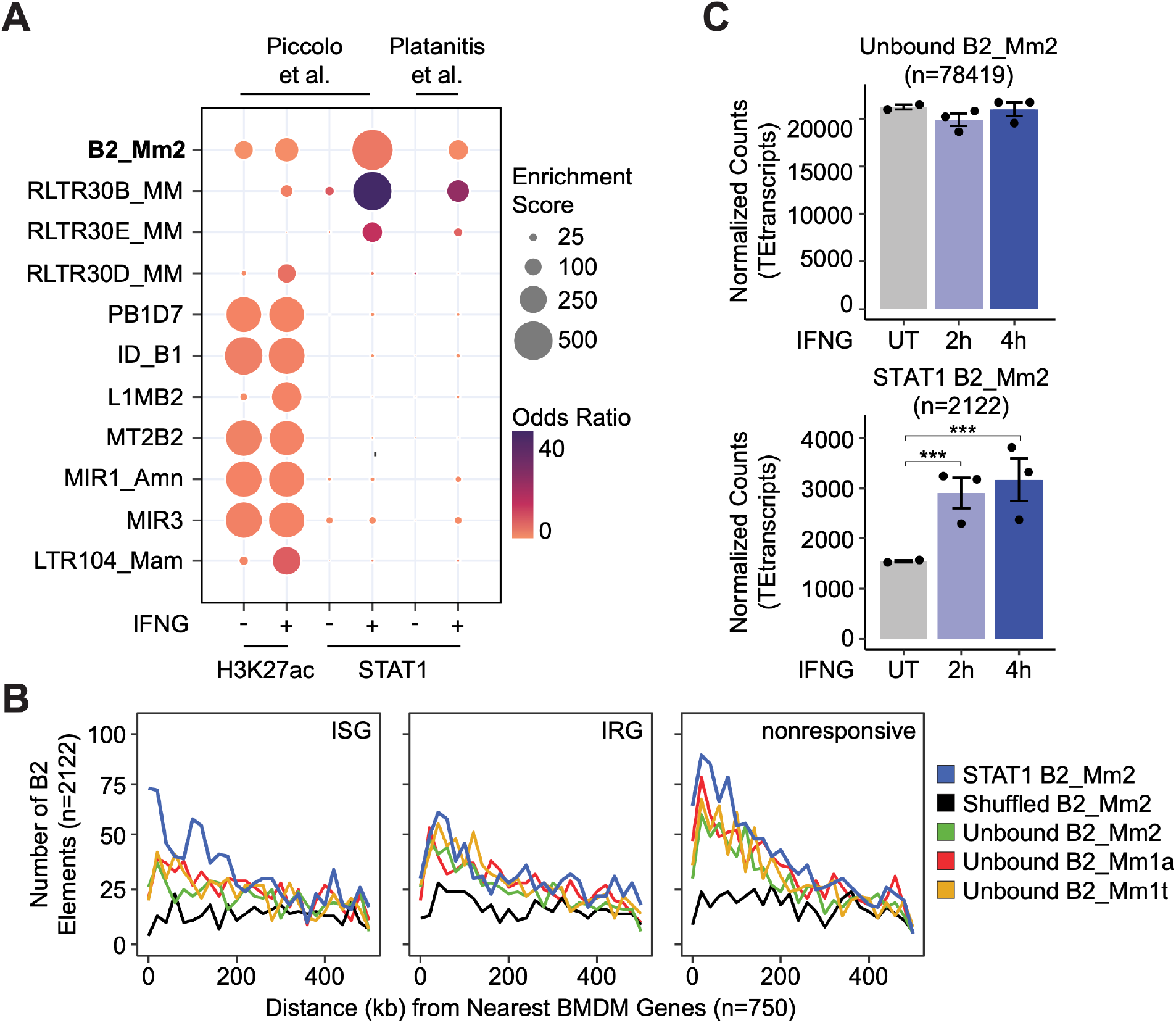
Identification of transposon-derived enhancers in innate immunity. (A) Bubble plot showing family-level enrichment of transposons within ChIP-seq peak regions. TE families are sorted by descending median Kimura distance. GIGGLE enrichment score is a composite of the product of both-log_10_(*p*-value) and log_2_(odds ratio). (B) Frequency histogram of absolute distances from STAT1-bound B2_Mm2 (blue, n=2122), randomly shuffled B2_Mm2 (black, n=2122), and randomly subset unbound B2_Mm2 (green, n=2122), B2_Mm1a (red, n=2122), and B2_Mm1t (yellow, n=2122) elements to the nearest ISG (n=750), IRG (n=750), or nonresponsive gene (n=750). Data shown for (32) comparing expression in BMDMs stimulated with IFNG for 4 hours relative to untreated. (C) DESeq2 normalized counts showing immune-stimulated expression of unbound B2_Mm2 (top, n=78419) and STAT1-bound B2_Mm2 (bottom, n=2122) elements in murine BMDMs. Data shown for untreated (n=2) BMDMs and BMDMs stimulated with IFNG for 2 hours (n=3) or 4 hours (n=3). Treatments are indicated by color. ***DESeq2 *p*-value < 0.0001. Error bars designate SEM. Data shown for (32). ISG: Interferon-stimulated gene; IRG: Interferon-repressed gene. BMDMs: Bone marrow derived macrophages. SEM: Standard error of mean.

We previously identified enrichment of RLTR30 elements within STAT1 binding sites in IFNG and IFNB-stimulated mouse macrophages based on analysis of a different ChIP-seq dataset (10, 23). However, our previous analysis did not capture enrichment of B2_Mm2, likely because the dataset was generated using 36 bp short reads. In contrast, the more recent datasets analyzed here used 50 bp reads (19), which improves mappability to individual copies of evolutionarily young TE families such as B2_Mm2 (24).

### B2_Mm2 elements contain STAT1 binding sites and show inducible enhancer activity

B2_Mm2 is a murine-specific subfamily of the B2 short interspersed nuclear element (SINE) family, which is highly abundant in the mouse genome. B2 SINE elements are divided into three subfamilies, including B2_Mm2 (80,541 copies), B2_Mm1a (16,321 copies), and B2_Mm1t (35,812 copies). B2 SINE elements have been characterized to show a wide range of regulatory activities in mice, including acting as promoters (25), insulator elements bound by CTCF (26–28), or regulatory non-coding RNAs (29–31). As the potential for B2_Mm2 SINEs to act as inducible enhancers has not yet been investigated, we decided to further investigate B2 SINEs in this context.

The B2_Mm2 subfamily showed strong evidence of enrichment within regions bound by STAT1, providing 2122 total binding sites (odds ratio = 4.85). These B2_Mm2 elements show significantly higher localization near ISGs (*p*-value = 5.03×10^-52^, odds ratio = 9.13) than interferon-repressed genes (IRGs) or nonresponsive genes, compared to unbound B2 elements or random genomic regions (Figure 1B, Supplemental Figure S1A-B). We did not observe consistent enrichment of the B2_Mm1a and B2_Mm1t subfamilies over STAT1-bound regions (Supplemental Table S4). Additionally, STAT1-bound B2_Mm2 elements are transcriptionally upregulated at the family level in response to IFNG stimulation (Figure 1C, Supplemental Figure S2A-C, Supplemental Table S5). Although unbound B2_Mm2 elements are also transcriptionally active, we did not observe a significant increase in expression in response to IFNG stimulation. Taken together, these data indicate that thousands of B2_Mm2 elements show epigenetic and transcriptional evidence of IFNG-inducible regulatory activity in primary murine bone marrow derived macrophages.

We investigated the sequence features of each B2 SINE subfamily to determine the basis of IFNG-inducible activity. Given that B2 SINE elements have previously been associated with CTCF binding due to the presence of a CTCF motif harbored by most copies (27), we subdivided elements from each family based on occupancy by STAT1, CTCF, both factors, or neither factor based on ChIP-seq. Across each of these subsets, we looked for the presence of GAS or CTCF motifs (Figure 2A, Supplemental Figure S3). As expected, all B2 subfamilies showed extensive ChIP-seq binding evidence of CTCF and the RAD21 cohesin subunit, coinciding with a CTCF motif (Supplemental Figure S3). In contrast, only a subset of elements from the B2_Mm2 subfamily showed inducible binding of STAT1 (Figure 2A). Consistent with ChIP-seq evidence, STAT1-bound B2_Mm2 elements contain both GAS and CTCF motifs, while B2_Mm1a/t elements only harbor CTCF motifs (Figure 2B, Supplemental Figure S4). Within B2_Mm2, elements that are bound by STAT1 are significantly enriched for GAS motifs when compared against unbound elements (E-value 2.08×10^-71^, Supplemental Table S6). In addition, STAT1-bound B2_Mm2 elements contain stronger sequence matches to GAS motifs compared to unbound B2_Mm2 elements and B2_Mm1a/t elements (Figure 2A, Supplemental Figure S5). Therefore, elements of the B2_Mm2 subfamily are uniquely characterized by GAS motifs that are associated with STAT1 binding activity.

**Figure 2.**
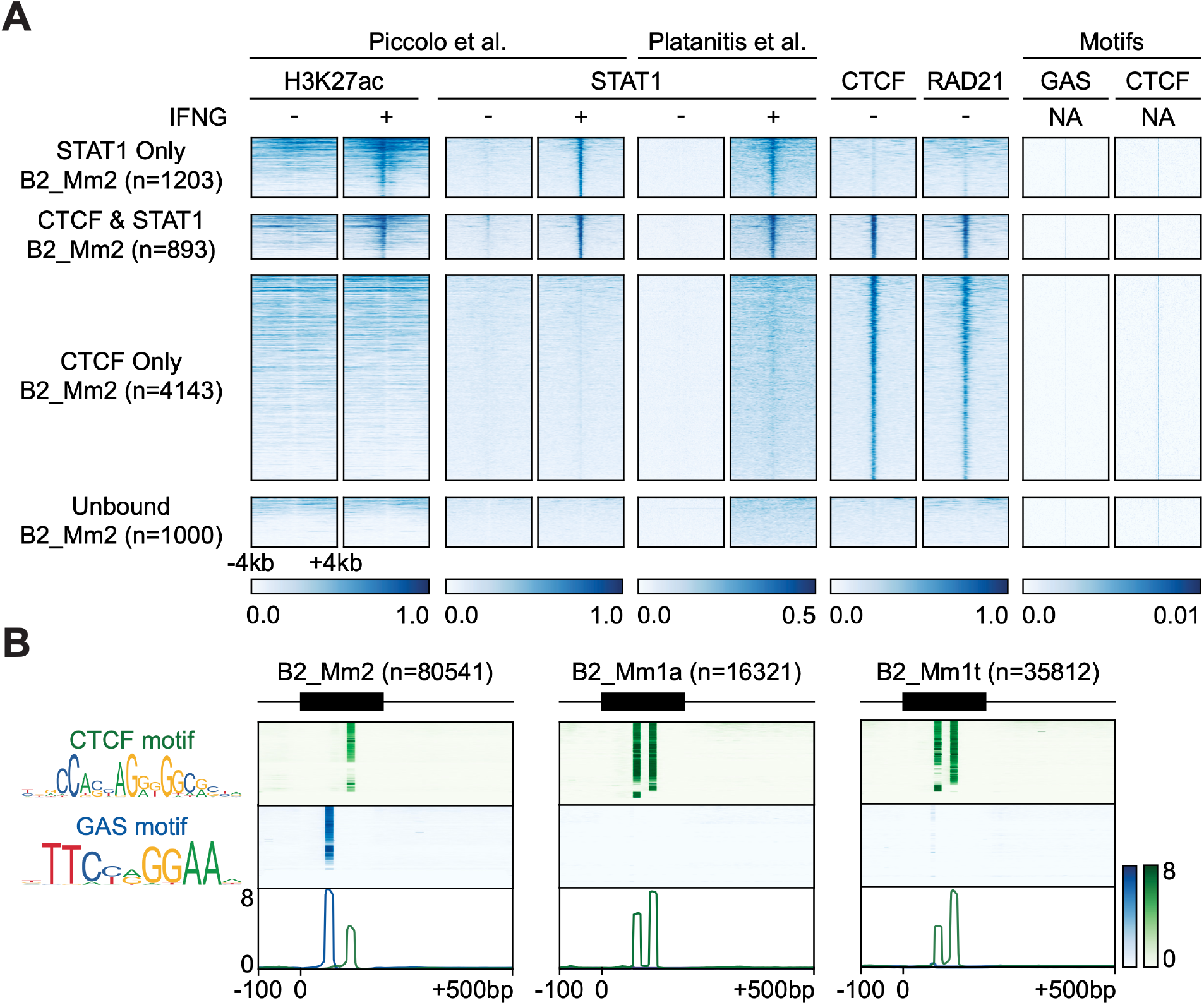
Epigenomic profiling of B2_Mm2. (A) Heatmaps showing CPM-normalized ChIP-seq signal and motif signal centered over B2_Mm2 elements bound only by STAT1 (n=1203); B2_Mm2 bound by both STAT1 and CTCF (n=893); B2_Mm2 bound only by CTCF (n=4143); and a random subset of unbound B2_Mm2 (n=1000). Regions are sorted by descending mean CPM signal. Signal intensity is indicated below. CTCF track derived from (48). RAD21 track derived from (63). (B) Schematic of GAS (blue) and CTCF (green) motifs present within extant B2_Mm2 (left, n=80541), B2_Mm1a (middle, n=16321), and B2_Mm1t (right, n=35812) sequences. Heatmap intensity corresponds to motif matches based on the log likelihood ratio. Heatmaps are sorted by descending mean signal. Position weight matrices were obtained from JASPAR (69). CPM: Counts per million. GAS: Gamma activated sequence.

Notably, we found that STAT1-bound B2_Mm2 elements are the only B2 elements that show inducible H3K27ac signal associated with enhancer activity. In contrast, B2_Mm2 elements bound only by CTCF or unbound elements show minimal H3K27ac signal (Figure 2A). B2_Mm1a/t elements also show minimal STAT1 signal regardless of CTCF binding (Supplemental Figure S3). This suggests that the binding of STAT1 to B2_Mm2 elements causes activation of enhancer activity including acetylation of H3K27. Thus, B2_Mm2 elements represent a distinct subclass of B2 SINE elements that exhibit IFNG-inducible enhancer activity.

### An intronic B2_Mm2 element functions as an inducible enhancer of Dicer1

Having established that B2_Mm2 elements are an abundant source of IFNG-inducible STAT1 binding sites in the mouse genome, we asked whether any of these elements have been co-opted to regulate expression of individual ISGs. Focusing on the Piccolo et al dataset (32), we identified 699 STAT1-bound B2_Mm2 elements that fall within 50 kb of an ISG (Supplemental Table S7). From this set, we discovered an element located on Chromosome 12 within the first intron of *Dicer1,* which is an endonuclease responsible for recognizing and cleaving foreign and double stranded RNA that has been linked to innate immunity (33–36). While the human ortholog *DICER1* does not show IFNG-inducible expression in human primary macrophages (10, 37), mouse *Dicer1* shows a significant 50% upregulation in response to IFNG in primary mouse BMDMs (Figure 3A). This indicates that *Dicer1* likely acquired IFNG-inducible expression in the mouse lineage, potentially due to the gain of a species-specific IFNG-inducible regulatory element. The intronic B2_Mm2 element (B2_Mm2. *Dicer1*) shows inducible STAT1 and H3K27ac signal as well as constitutive binding by CTCF and RAD21 (Figure 3B). The element provides the only prominent nearby STAT1 binding site and is not present in rat or other mammals (Figure 3B). Therefore, we hypothesized that the B2_Mm2. *Dicer1* element was co-opted as an IFNG-inducible enhancer of mouse *Dicer1*.

**Figure 3.**
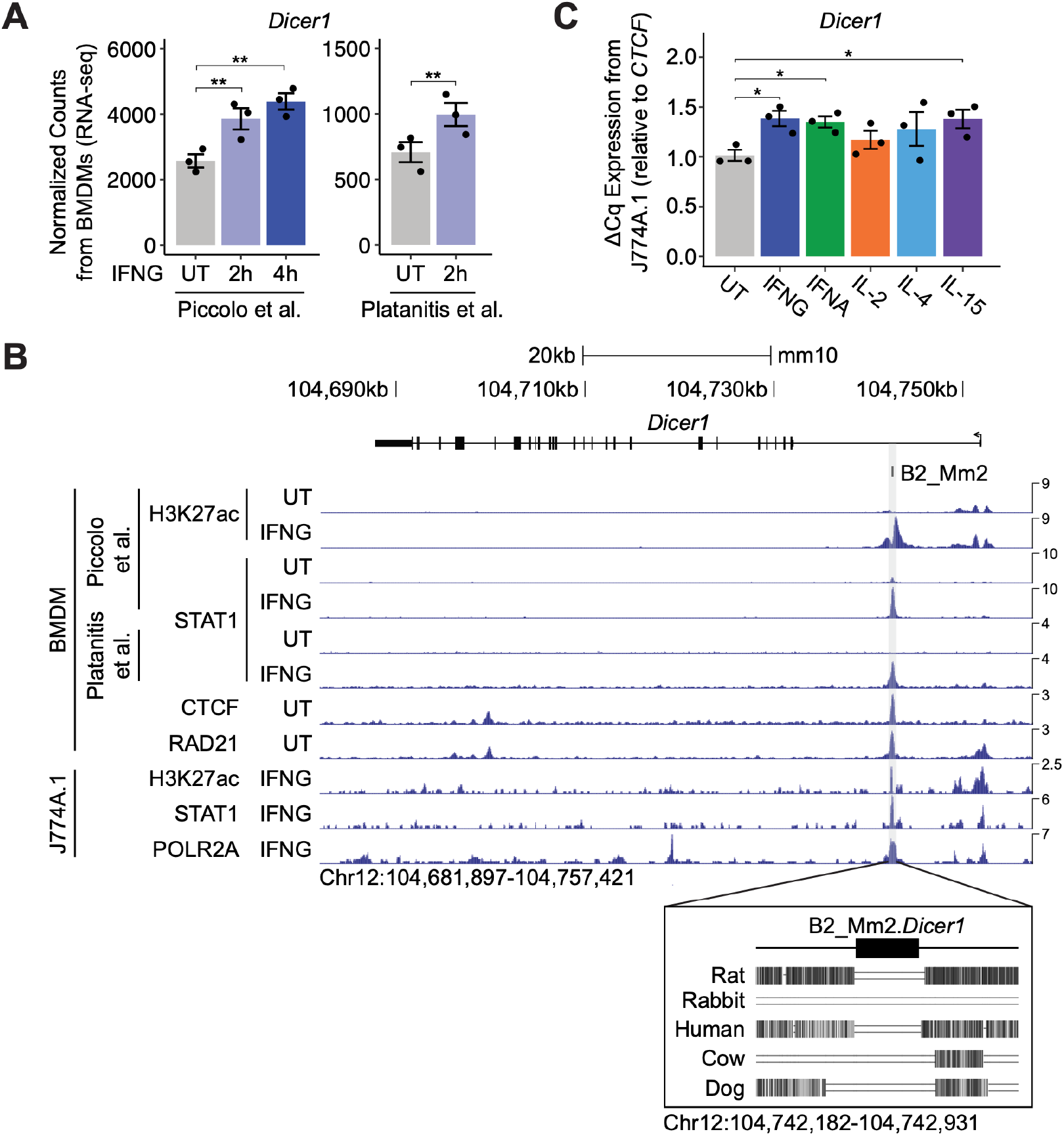
Identification of a putative B2_Mm2-derived enhancer for Dicer1. (A) DESeq2 normalized counts showing immune-stimulated expression of *Dicer1* in primary murine BMDMs. Data shown for untreated (n=3) BMDMs and BMDMs stimulated with IFNG for 2 hours (n=3) or 4 hours (n=3). **DESeq2 *p*-value < 0.001. Error bars designate SEM. (B) Genome browser screenshot (http://genome.ucsc.edu) of the *Dicer1* locus (Chr12:104,681,897-104,757,421) showing CPM-normalized ChIP-seq tracks for primary murine BMDMs and immortalized macrophage line J774A.1. B2_Mm2.*Dicer1* (Chr12:104,742,467-104,742,646) is highlighted in grey. Values on the right of each track correspond to signal maxima. Bottom inset shows B2_Mm2.*Dicer1* with accompanying conservation tracks for rat, rabbit, human, cow, and dog. CTCF track derived from (48). RAD21 track derived from (63). (C) RT-qPCR of wild type untreated (grey, n=3) J774A.1 cells and J774A.1 cells stimulated with IFNG (blue, n=3), IFNA (green, n=3), IL-2 (orange, n=3), IL-4 (light blue, n=3), or IL-15 (purple, n=3) for 4 hours. Treatments are indicated by color. \**p*-value < 0.05, Student’s paired two-tailed *t*-test. BMDM: Bone marrow derived macrophage. SEM: Standard error of mean. CPM: Counts per million.

To experimentally test the potential enhancer activity of B2_Mm2.*Dicer1*, we used the mouse J774A.1 macrophage-like cell line, a commonly used model of murine immunity (38, 39). We first confirmed using RT-qPCR that *Dicer1* shows 30-40% upregulation after 4 hrs of IFNG treatment (Figure 3C). Given the STAT1 binding site and motif present in B2_Mm2.*Dicer1*, we also tested other cytokines that act through STAT-family transcription factors. We found that 4 hr treatment of J774A.1 cells with IFNA, IL6, and IL4 all induced *Dicer1* expression to similar levels (30-40%), consistent with inducible regulation of *Dicer1* by JAK-STAT signaling, potentially through the STAT binding site present in B2_Mm2.*Dicer1*.

We next used CRISPR to generate clonal J774A.1 lines harboring homozygous deletions of the B2_Mm2.*Dicer1* element. We delivered guide RNAs targeting the flanking boundaries of B2_Mm2.*Dicer1* along with recombinant Cas9 by electroporation (Supplemental Figure S6A-C), and screened clonal lines for homozygous deletions by PCR (Supp fig. 6D-F). We isolated 2 clonal cell lines with a homozygous knockout of B2_Mm2.*Dicer1*, along with multiple wild-type (WT) J774A.1 clonal lines that were not electroporated to control for potential effects of clonal expansion (Supplemental Figure S7A). We used RT-qPCR to compare *Dicer1* expression levels and inducibility by IFNG in knockout and WT clones. WT clones showed consistent inducible expression, while both knockout clonal lines showed a complete lack of inducible expression (Figure 4A). These experiments demonstrate that B2_Mm2.*Dicer1* acts as an IFNG-inducible enhancer of *Dicer1* in J774A.1 cells.

**Figure 4.**
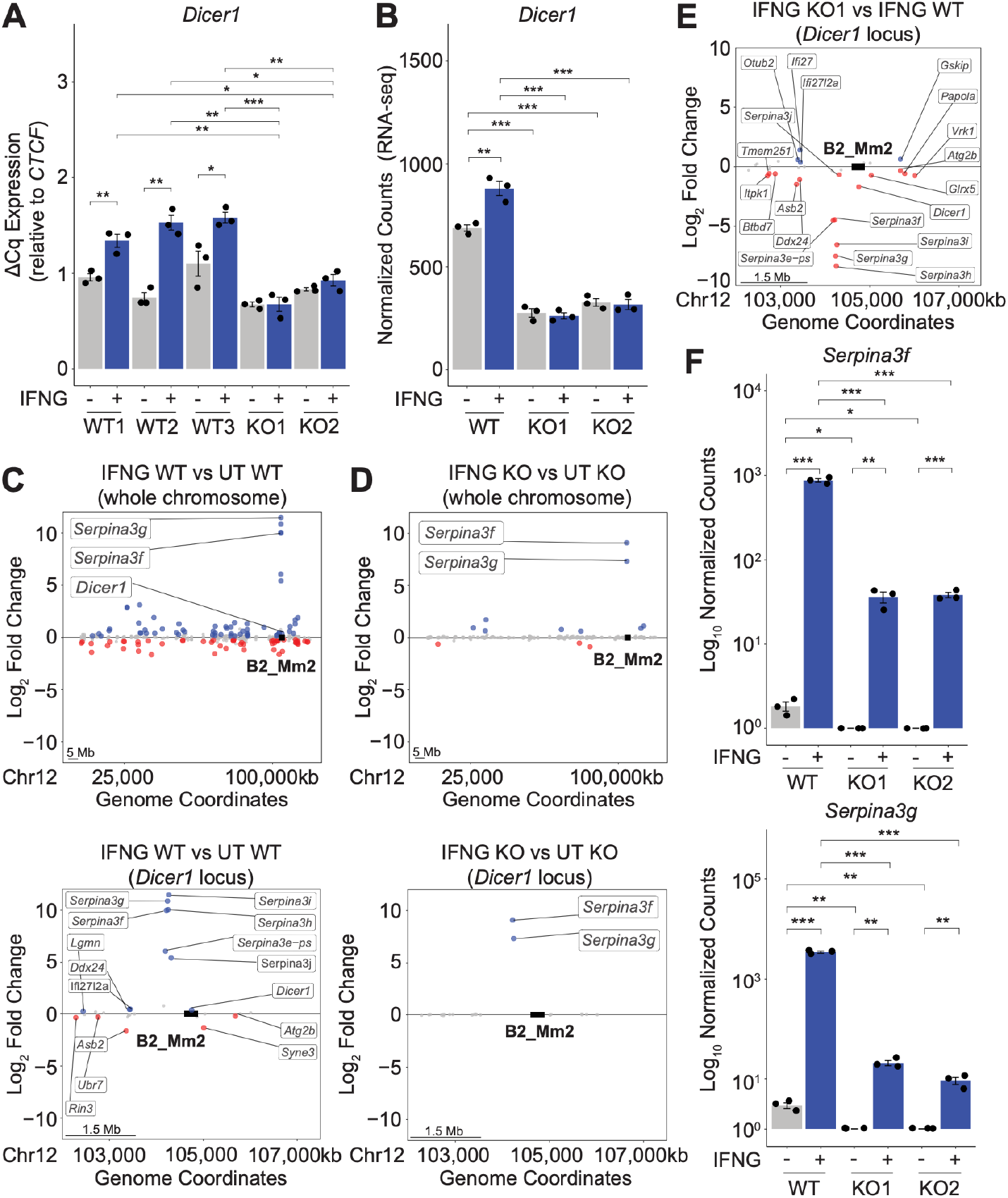
B2_Mm2 in the genomic landscape. (A) RT-qPCR of clonal replicates for WT (n=3) J774A.1 cells and J774A.1 cells harboring the B2_Mm2.Dicer1 deletion (n=2) measuring *Dicer1* expression relative to *CTCF*. Treatments are indicated by color. \**p*-value < 0.05, \*\**p*-value < 0.01, \*\*\**p*-value < 0.001, Student’s paired twotailed *t*-test. (B) DESeq2 normalized counts of *Dicer1* expression in bulk WT and knockout B2_Mm2.*Dicer1* J774A.1 cells. Treatments are indicated by color. \**p*-value < 0.05, \*\**p*-value < 0.01, \*\*\**p*-value < 0.001. (C) Distance plot visualizing changes in gene expression in wild type J774A.1 cells in response to IFNG for Chr12 (top) and over a 5Mb window centered on B2_Mm2.*Dicer1* (bottom). Significantly downregulated (log_2_FC < 0, FDR < 0.05) genes are shown in red while significantly upregulated (log_2_FC > 0, FDR < 0.05) genes are shown in blue. B2_Mm2.*Dicer1* is represented as a black box (not drawn to scale). (D) Same as in (C) but visualizing changes in gene expression in KO J774A.1 cells in response to IFNG. (E) Same as in (C) but visualization differences in gene expression in B2_Mm2.*Dicer1* KO cells relative to WT in response to IFNG. (F) DESeq2 normalized counts for *Serpina3f* and *Serpina3g* in WT and B2_Mm2.*Dicer1* KO J774A.1 cells. Treatments are indicated by color. \**p*-value < 0.05, \*\**p*-value < 0.01, \*\*\**p*-value < 0.001, Student’s paired two-tailed *t*-test.

### B2_Mm2.Dicer1 impact on the genomic regulatory landscape

We used RNA-seq to study the genome-wide effects of the B2_Mm2.*Dicer1* element in both knockout clones and a WT line. Consistent with the RT-qPCR results, we found that *Dicer1* showed significant 30% upregulation in WT cells but that this induction was completely ablated in knockout B2_Mm2 cells (Figure 4B). Notably, the RNA-seq normalized count data revealed that levels of *Dicer1* were also significantly reduced in untreated knockout cells (Figure 4B, 5A). This indicates that the B2_Mm2.*Dicer1* element also regulates *Dicer1* basal expression, potentially mediated by the constitutive CTCF binding site also contained within the element. By plotting IFNG-inducible gene expression in both WT and knockout clones across Chromosome 12, we found that deletion of the B2_Mm2.*Dicer1* element led to loss or repression of IFNG-inducible expression of multiple genes nearby the deletion (Figure 4C-E, Supplemental Figure 7B-E).

**Figure 5.**
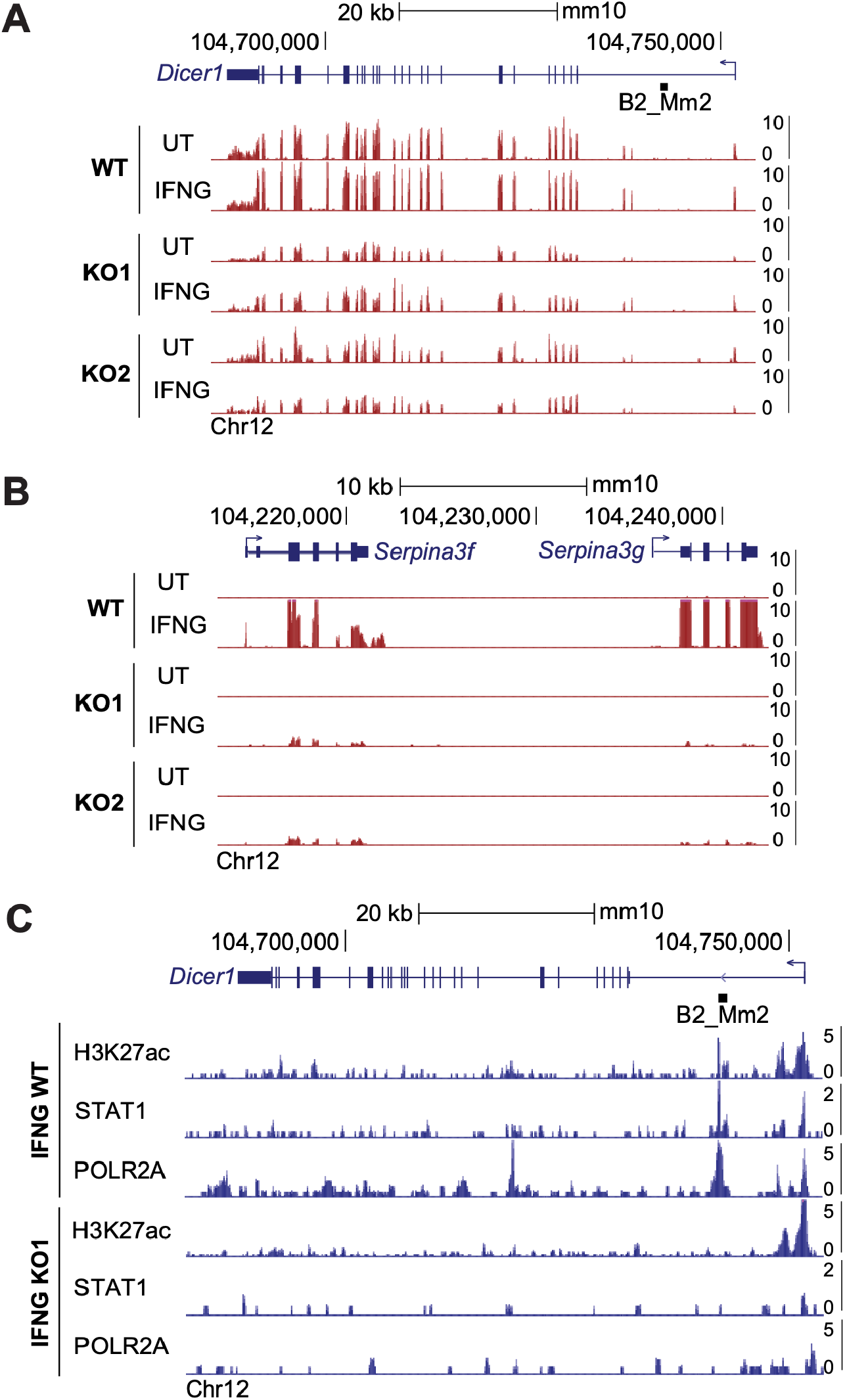
B2_Mm2 impacts local chromatin profile. (A) Genome browser screenshot (http://genome.ucsc.edu) of the *Dicer1* locus visualizing CPM-normalized expression in WT and B2_Mm2.*Dicer1* KO J774A.1 cells. Values on the right of each track correspond to signal maxima. B2_Mm2.Dicer1 is represented as a black box (not drawn to scale). (B) Same as in (A) but visualizing the *Serpina3f* and *Serpina3g* loci. (C) Genome browser screenshot of the *Dicer1* locus showing CUT&TAG data from WT and B2_Mm2.*Dicer1* KO J774A.1 cells.

Our RNA-seq analysis revealed that deletion of B2_Mm2.*Dicer1* also had a significant repressive effect on the IFNG-inducible expression of *Serpina* genes. In mouse, *Serpina* is a family of inflammatory genes represented by a large cluster of 14 murine paralogs, located 330 - 1000 kb from B2_Mm2.*Dicer1*. In particular, we found that *Serpina3f* and *Serpina3g* (approximately 500 kb from B2_Mm2.*Dicer1*) show a 1-2 orders of magnitude reduction in basal and induced gene expression in knockout cells compared to WT cells (Figure 4E-F, 5B). The relatively long range of this regulatory activity is not uncommon in vertebrate genomes (40) and is potentially mediated by the CTCF binding activity of the element.

Finally, we confirmed the absence of intronic enhancer activity in B2_Mm2.*Dicer1* knockout cells. We used CUT&TAG (41) to profile H3K27ac, phosphorylated RNA polymerase II subunit A (POLR2A), and STAT1. In IFNG-stimulated WT cells, the B2_Mm2.*Dicer1* element shows prominent H3K27ac, STAT1, and POLR2A signal. However, these signals are completely lost in the knockout clone (Figure 5C). Collectively, these experiments confirm that B2_Mm2.Dicer1 has been co-opted to function as an IFNG-inducible enhancer that regulates multiple immune-related genes in the *Serpina-Dicer1* locus.

## DISCUSSION

B2 SINE elements are abundant in the mouse genome and they have been widely studied due to their substantial influence on genome regulation and evolution. B2 elements have contributed non-coding RNAs inducible by stress or infection (29, 42–46), splicing signals (47), promoter elements (25), and CTCF-bound insulator elements (26–28). Our study reveals a new subclass of B2 elements that have IFNG-inducible enhancer activity. These elements, which belong to the B2_Mm2 subfamily, contain strong binding sites for both STAT1 and CTCF, and have the potential to exert long-range enhancer regulation.

Given the abundance of B2 elements and their potential to cause pathological regulatory rewiring, many B2 elements are targeted for SETDB1/H3K9me3-mediated epigenetic repression which inhibits their regulatory potential (48). Therefore, the functional impact of B2 elements on the mouse epigenome remains unclear. By using CRISPR to generate knockout cells of a B2_Mm2 element, we demonstrated that B2 elements can be co-opted to act as inducible enhancer elements in the context of IFNG stimulation. While our experiments were conducted in the J774.A1 immortalized cell line, we confirmed that thousands of B2_Mm2 elements including B2_Mm2. *Dicer1* show strong transcriptional and epigenetic signatures of inducible enhancer activity in multiple primary macrophage epigenomic datasets.

Our identification of a B2_Mm2 element as an intronic enhancer of *Dicer1* potentially uncovers a novel regulatory feedback loop that controls *Dicer1* function related to TE silencing. Previous studies have demonstrated that *Dicer1* cleaves double-stranded RNAs including those derived from B2 SINE transcripts (49). In *Dicer1* knockout embryonic stem cells, TE-derived transcripts are upregulated (50), and upregulation of B2-derived double-stranded RNAs causes activation of the IFN response (36). Therefore, *Dicer1* is important for defense against aberrant TE upregulation. We speculate that co-option of the B2_Mm2 element as an enhancer of *Dicer1* facilitates upregulation of *Dicer1* in response to conditions that drive TE upregulation, such as infection or stress.

In human, dysregulation of *DICER1* is associated with a wide range of pathologies ranging from *DICER1* syndrome, cancer, neurological diseases such as Parkinson’s disease, and autoimmune disorders such as rheumatoid arthritis (51). While *DICER1* has a highly conserved function as an endonuclease involved in RNA interference, our work highlights that orthologs of *DICER1* has undergone lineage- or species-specific regulatory evolution that may drive underappreciated differences in function across species. This could have significant implications when developing and testing RNA-based therapeutics in mouse genetic models, which may elicit distinct *Dicer1*-mediated responses due to species-specific enhancers such as B2_Mm2.*Dicer1*.

Finally, this work adds to a growing list of studies demonstrating the co-option of lineagespecific TEs to regulate ISGs. Previous genomic and experimental work in human (10) and cow cells (52) have revealed independent co-option of TEs as IFNG-inducible enhancer elements. Notably, while endogenous retroviruses are the major source of TE-derived inducible enhancers in human, they have a relatively minor contribution in cow and mouse. Instead, we found that the B2_Mm2 SINE subfamily is the predominant source of TE-derived inducible enhancers in mouse, and the Bov-A2 SINE subfamily is the predominant source in cow (52). These findings suggest that the acquisition of STAT1-associated GAS motifs and enhancer activity is not restricted to any one type of TE. It remains unclear whether the IFNG-inducible regulatory activity promotes TE replication, or whether the emergence of these motifs is fortuitous and has no effect on TE fitness. Nonetheless, our work supports the idea that TEs have been repeatedly co-opted to as IFNG-inducible enhancers throughout mammalian evolution and contribute to the divergence of immune regulatory networks.

## METHODOLOGY

### Sequences

A list of all primer sequences and gRNA sequences can be found in Supplemental Table S8.

### RNA-seq Analysis

Single-end RNA-seq data from primary murine BMDMs stimulated with 100 ng/mL IFNG for 2 or 4 hours (32) or 10 ng/mL IFNG for 2 hours (19) were downloaded from SRA using fasterq-dump v3.0.0 (53). Adapters and low-quality reads were trimmed using BBDuk v38.05 (54) using options ‘*ktrim*=*r k*=*34 mink*=*11 hdist*=*1 qtrim*=*r trimq*=*10 tpe tbo*’. Library quality was assessed using FastQC v0.11.8 (55) and MultiQC v1.7 (56), and trimmed reads were aligned to the mm10 assembly using HISAT2 v2.1.0 (57) with option *‘--no-softclip’*. Only uniquely aligned fragments (MAPQ >= 10) were retained using samtools v1.10 (58). Aligned fragments were assigned to the complete mm10 Gencode M18 (59) annotation in an unstranded manner using featureCounts v1.6.2 (60) with options *‘-p -O -s 0 -t exon -g gene_id’*, and differentially expressed genes between IFNG-stimulated and unstimulated cells were called using DESeq2 v1.26.0 (61). For most analyses, ISGs and IRGs were defined as genes with a false discovery rate (FDR) of at least 0.05 and log_2_ fold change (log_2_FC) greater than 0 and less than zero, respectively. Nonresponsive genes were defined using the following cutoffs: baseMean greater than 100; FDR greater than 0.90; and absolute log2FC less than 0.10. Interferon stimulation was confirmed by gene ontology analysis using gProfiler (last updated 05/18/2022) with FDR < 0.05 (62).

### ChIP-seq Analysis

Single-end ChIP-seq data from primary murine BMDMs (19, 32, 48, 63) were downloaded from SRA using fasterq-dump v3.0.0 (53). Adapters and low-quality reads were trimmed using BBDuk v38.05 (54) using options ‘*ktrim*=*r k*=*34 mink*=*11 hdist*=*1 qtrim*=*r trimq*=*10 tpe tbo*’. Library quality was assessed using FastQC v0.11.8 (55) and MultiQC v1.7 (56), and trimmed reads were aligned to the mm10 assembly using BWA-MEM v0.7.15 (64). Low quality and unmapped reads were filtered using samtools v1.10 (58), and duplicates were removed with

Picard MarkDuplicates v2.6.0 (65). Peak calling was performed with MACS2 v2.1.1 (66) using options ‘--*gsize mm* –*pvalue 0.01* –*bdg* –*SPMR* –*call-summit*’. bigWigs corresponding to read pileup per million reads for visualization on the UCSC Genome Browser (67). Where possible, only peaks overlapping more than one replicate were retained for further analysis. To confirm whether STAT1 peaks were enriched for their associated binding motifs, we ran XSTREME v5.4.1 (68) using options ‘--*minw 6* – *maze 20* –*streme-nmotifs 20* –*align center*’ querying against the JASPAR CORE 2018 vertebrates database (69).

### Transposable Element Analysis

To identify TE families enriched for STAT1 peaks, we used GIGGLE v0.6.3 (22) to create a database of all TE families annotated in the mm10 genome according to Dfam v2.0 (70) annotation. ChIP-seq peaks were then queried against each TE family in the database. We only retained TE families with a reported GIGGLE enrichment score greater than one for both H3K27ac and STAT1. Results were visualized as a bubble plot where the filtered TE families were sorted by ascending Kimura divergence according to RepeatMasker (71) output. Reported odds ratios and *p*-values are derived from Fisher’s exact test. For further analysis, we intersected STAT1 peaks with the full TE annotation or B2 elements specifically using BEDTools v2.28.0 (72). For the heatmap visualizations using deepTools v3.0.1 (73), signal from counts per million- (CPM) normalized bigWigs was plotted over a subset of B2_Mm2 elements that are bound only by STAT1, CTCF, both, or neither. We additionally visualized ChIP-seq signal over all B2_Mm2, B2_Mm1a, and B2_Mm1t elements by descending average signal, excluding elements with zero overlapping signal.

To assess whether STAT1-bound B2_Mm2 elements are enriched near ISGs, we sorted all ISGs and IRGs by descending and ascending log_2_FC, respectively, and retained the top 750 genes. We additionally randomly subset for 750 nonresponsive genes. The absolute distance to the nearest ISG, IRG, or nonresponsive gene was determined for all STAT1-bound B2_Mm2 elements using BEDTools v2.28.0 (72). Randomly shuffled STAT1-bound B2_Mm2 as well as randomly subset, unbound B2_Mm2, B2_Mm1a, and B2_Mm1t were included as controls. Statistical significance was determined for the first 20kb bin using the Fisher’s exact test with BEDTools v2.28.0 (72).

To identify TE families that are differentially expressed in response to IFNG in primary murine BMDMs, we realigned the RNA-seq data to the mm10 reference genome using HISAT2 v2.1.0 (57) with options ‘-*k100* –*no-softclip*’. Aligned reads were assigned to TE families using TEtranscripts v2.1.4 (74) with options ‘--*sortByPos* –*mode multi* –*iteration 100* –*stranded no*’. To differentiate unbound and STAT1-bound B2_Mm2 elements, we generated a custom TE annotation file compatible with TEtranscripts that includes all TEs annotated in Dfam v2.0 (70) but annotates STAT1-bound B2_Mm2 elements as an independent subfamily. Differentially expressed TE families between IFNG-stimulated and unstimulated cells were identified using DESeq2 v1.26.0 (61). TE families with an FDR less than 0.05 and log_2_FC greater than 0.50 were considered as differentially expressed.

We identified putative STAT1 and CTCF binding sites genome-wide using FIMO v5.0.3 (75) with a *p*-value cutoff of 1×10^-4^ (heatmaps) or 1 (B2 box-and-whisker). For all motif analyses, binding motif position-weight matrices for STAT1 and CTCF were obtained from the JASPAR CORE 2018 vertebrate database (69). To visualize motif presence over all B2 elements, repeat 5’ start coordinates were recalculated based on their alignment to the consensus according to RepeatMasker annotations. Motif presence was visualized as a heatmap using deepTools v3.0.1 (73), and elements were sorted by descending average signal. We additionally aligned the consensus sequences for B2_Mm2, B2_Mm1a, and B2_Mm1t from Repbase v24.02 and the sequence for B2_Mm2.*Dicer1* using MUSCLE v3.8.1551 (76). Predicted STAT1 and CTCF motifs were identified using FIMO v.5.0.3 (75), and base changes relative to the canonical binding motifs were highlighted according to the weight of each individual base in the position-weight matrices. Finally, we filtered for STAT1-bound B2_Mm2 elements that were nonoverlapping, non-nested, and unique and ran AME v5.4.1 (77) using a subset of unbound B2_Mm2 elements as the background control with options ‘--*kmer 2* –*method fisher* –*scoring avg*’.

### Cell Line Passing and Interferon Treatments

J774A.1 mouse cells (ATCC) were cultured in DMEM supplemented with 1X penicillinstreptomycin and 10% fetal bovine serum. J774A.1 cells were routinely passaged using 0.25% Trypsin-EDTA and cultured at 37°C and 5% CO2. All IFNG treatments were performed using 100 ng/mL recombinant mouse IFNG (R&D Systems #485-MI-100).

### Cytokine Panel

All treatments were carried out using a 4 hour time period. IL-4 (Sigma-Aldrich #I1020-5UG) was added to a final concentration of 1 ng/mL, recombinant IFNA (R&D Systems #12100-1) to a final concentration of 1000 U/mL, recombinant IL-2 (R&D Systems #402-ML-020) to a final concentration of 100 ng/mL, and recombinant IL-6 (Sigma-Aldrich #I9646-5UG) to a final concentration of 20 ng/mL in accordance with manufacturers’ recommendations. RNA was extracted using an Omega Mag-Bind Total RNA Kit (Omega Bio-Tek #M6731-00) and analyzed via RT-qPCR.

### Design of gRNA constructs

Two gRNA sequences were designed to flank each side of B2_Mm2.*Dicer1* in order to delete the element and generate knockout J774A.1 cells via SpCas9 (Integrated DNA Technologies #1081060). All gRNA sequences were also verified to uniquely target the locus of interest using the UCSC BLAT tool (78) against the mm10 genome assembly.

### Generation of CRISPR KO Cell Lines

After gRNAs were designed, we used Alt-R Neon electroporation (1400V pulse voltage, 10msec pulse width, 3 pulses total) with four different combinations of the gRNAs to target B2_Mm2. One set of guides was found to produce the expected doubles-stranded cuts on both sides of the element in the bulk electroporated cell populating using gel electrophoresis. Clonal lines were isolated using the array dilution method and screened for the expected homozygous deletion using primers flanking the B2_Mm2.*Dicer1* element. Clonal lines homozygous for the deletion were further validated using one flanking primer and one primer internal to B2_Mm2.*Dicer1*. To determine deletion breakpoint sequences, PCR products flanking each deletion site were cloned into a sequencing vector using the CloneJET PCR Cloning Kit (Thermo Fisher Scientific #K1231) and transformed into 5-alpha Competent *E. coli* (New England Biolabs #C2987H). Plasmid DNA was harvested using the EZNA Omega Plasmid DNA Mini Kit I (Omega Bio-Tek #D6942-02), and the sequence of each construct was verified by Sanger sequencing (Quintara Biosciences, Fort Collins, CO; Genewiz, South Plainfield, NJ). Sequencing results were visualized by aligning to the mm10 reference genome using BLAT (78). We identified two clonal lines homozygous for the B2_Mm2.*Dicer1* deletion for further experimentation.

### Quantifying *Dicer1* Expression using RT-qPCR

Real time quantitative polymerase chain reaction (RT-qPCR) was used to quantify *Dicer1* expression in WT and knockout cell lines. WT J774A.1 cells used in RT-qPCR are all biological replicates that underwent the same single cell seeding process as the KOs to serve as a control. Forward and reverse primers were designed for *CTCF, Dicer1, Gbp2b, Serpina3f*, and *Serpina3g*. Each set of primers was designed using a combination of tools from NCBI Primer BLAST (79), Benchling (Benchling), and IDT RT-qPCR Primer Design. The final primer sequences chosen were confirmed to uniquely bind to the desired target sequence using BLAT (78). RT-qPCR reactions were prepared using the Luna Universal One-Step RT-qPCR Kit (New England Biolabs #E3005S) according to the manufacturer’s instructions. RT-qPCR data were analyzed using *CTCF* as a housekeeping gene. A Cq, deltaCq, and deltaDeltaCq value were obtained for each well. These values were averaged to arrive at a mean deltaDeltaCq expression value for each treatment and genotype condition. Standard deviation and a twotailed Student’s *t*-test were then calculated for each treatment and genotype condition. A *p*-value of less than 0.05 demonstrates there is a statistically significant difference in gene expression levels between two treatment and/or genotype conditions.

### J774A.1 RNA-seq Library Preparation

The Zymo Quick RNA Miniprep Plus Kit (Zymo Research #R1504) was used to extract RNA from J774A.1 cells for all treatments except for the cytokine panel which used the Omega RNA Extraction Kit (Omega Bio-Tek #M6731-00). WT J774A.1 cells used in RNA-seq and all downstream analysis were a bulk population of J774A.1 cells that did not undergo single cell seeding. All RNA lysates and single-use aliquots of extracted RNA were stored at −80°C until library preparation. RNA integrity was quantified with High Sensitivity RNA TapeStation 4200 (Agilent). Libraries were generated using the KAPA mRNA HyperPrep Kit (KAPA Biosystems #08098123702) according to the manufacturer’s protocol. The final libraries were pooled and sequenced on a NovaSeq 6000 as 150bp paired-end reads (University of Colorado Genomics Core).

### J774A.1 RNA-seq Analysis

Adapters and low quality reads were first trimmed using BBDuk v38.05 (54). Library quality was assessed using FastQC v0.11.8 (55) and MultiQC v1.7 (56) and trimmed reads were aligned to the mm10 assembly using HISAT2 v2.1.0 (57) with option ‘--*rna-strandness RF*’. Only uniquely aligned fragments were retained, and technical replicates were merged using samtools v1.10 (58). CPM-normalized, stranded bigWigs were generated using deepTools bamCoverage v3.0.1 (73) and visualized using the UCSC Genome Browser (67). Aligned fragments were assigned to the mm10 refseq gene annotation in a reversely stranded manner using featureCounts v1.6.2 (60) with options ‘-*t exon -s 2*’, and differentially expressed genes were called using DESeq2 v1.26.0 (73). We analyzed every individual pairwise comparison to determine the effects due to both treatment and genotype. Untreated and wild type conditions were defined as the reference level. Log_2_FC values were shrunken using the apeglm function v1.8.0 (80) for visualization across Chromosome 12 as a distance plot.

### CUT&Tag

CUT&Tag chromatin pulldowns were generated using a protocol from (41, 81) with an input of 100-500k cells and the following modifications: pAG-Tn5 (EpiCypher #15-1017) was diluted 1:40 in nuclease-free water containing 20mM HEPES pH 7.5, 300mM NaCl, 0.5mM spermidine, 0.01% digitonin, and a protease inhibitor tablet, and libraries were amplified for 14 cycles. The following primary antibodies were used: rabbit IgG (1:1000, EpiCypher #13-0042), rabbit anti-H3K27ac (1:100), rabbit anti-pRPB1-Ser5 (1:100, Cell Signaling Technology #13523S), rabbit anti-STAT1 (1:100, Cohesion Biosciences #CPA3322), rabbit anti-pSTAT1-Ser727 (1:100, Active Motif #39634), rabbit anti-CTCF (1:100, EMD Millipore #07-729-25UL). Guinea pig antirabbit IgG (1:100, Antibodies-Online #ABIN101961) was used as a secondary antibody. CUTANA pAG-Tn5 (EpiCypher #15-1017) was added to each sample following primary and secondary antibody incubation. Pulldown success was measured by Qubit dsDNA High Sensitivity (Invitrogen) and TapeStation 4200 HSD5000 (Agilent) before proceeding to library preparation. Pulldowns were concentrated and pooled using KAPA Pure Beads (Roche). The final pooled libraries were quantities with TapeStation 4200 HSD5000 and sequences on an Illumina NovaSeq 6000 as 150bp paired-end reads (University of Colorado Genomics Core).

### CUT&Tag Analysis

Adapters and low quality reads were first trimmed using BBDuk v38.05 (54). Library quality was assessed using FastQC v0.11.8 (55) and MultiQC v1.7 (56). Trimmed reads were then aligned to the mm10 assembly using BWA-MEM v0.7.15 (64) and samtools (58) retained only uniquely aligned fragments (MAPQ >= 10). Peaks were called without a control file using MACS2 v2.1.1 (66). bigWigs corresponding to read pileup per million reads for visualization on the UCSC Genome Browser (67).

### External Datasets

Publicly available data were downloaded from public repositories using fasterq-dump from the NCBI SRA Toolkit. RNA-seq datasets were obtained from GSE84517 and GSE115434. ChIP-seq datasets were obtained from GSE84518, GSE108805, GSE189971, and GSE115433.

### Data Access

Raw and processed sequencing data generated in this study have been submitted to the NCBI Gene Expression Omnibus (GEO) with accession number GSE202574.

## Code Availability

UCSC Genome browser sessions and all code available at https://genome.ucsc.edu/s/coke6162/B2_SINE_enhancers and https://github.com/coke6162/B2_SINE_enhancers.

## Authors’ Contributions

EC, IH, and CK designed the study. IH, CK, and DS performed experiments. EC, IH, and CK analyzed data and interpreted results. IH and CK created and edited figures and tables. EC, IH, and CK wrote the manuscript with input from all co-authors. All authors gave final approval for publication.

## Acknowledgements

We thank the University of Colorado Genomics Shared Resource and BioFrontiers Computing core for technical support during this study.

