## Supplementary Info for "Mouse B2 SINE elements function as IFN-inducible enhancers"

### Supporting Information for
Mouse B2 SINE elements function as IFN-inducible enhancers.

Isabella Horton*, Conor J. Kelly*, David M. Simpson, Edward B. Chuong
*Equal contributions

Edward B. Chuong


**This PDF file includes:**

Figures S1 to S7
Legends for Tables S1 to S8
SI References

**Other supporting materials for this manuscript include the following:**

Tables S1 to S8

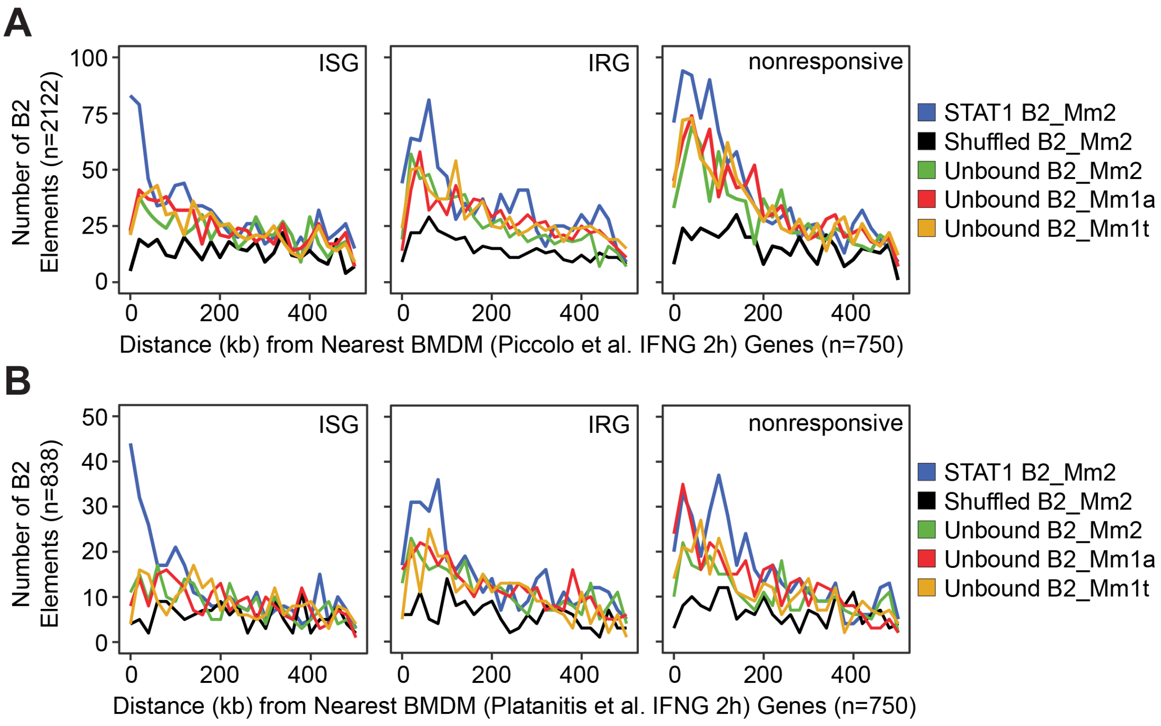


**Fig S1.** Distances from each B2 element to the nearest BMDM ISG. (A) Frequency histogram of absolute distances from STAT1-bound B2_Mm2 (blue, n=2122), randomly shuffled B2_Mm2 (black, n=2122), and randomly subset unbound B2_Mm2 (green, n=2122), B2_Mm1a (red, n=2122), and B2_Mm1t (yellow, n=2122) elements to the nearest ISG (n=750), IRG (n=750), or nonresponsive gene (n=750). Data shown for Piccolo et al. (1) where BMDMs were stimulated with IFNG for 2 hours. (B) Same as in (A) but for a subset of B2 elements (n=838) marked by STAT1. Data shown for Platanitis et al. (2) where BMDMs were stimulated with IFNG for 2 hours.

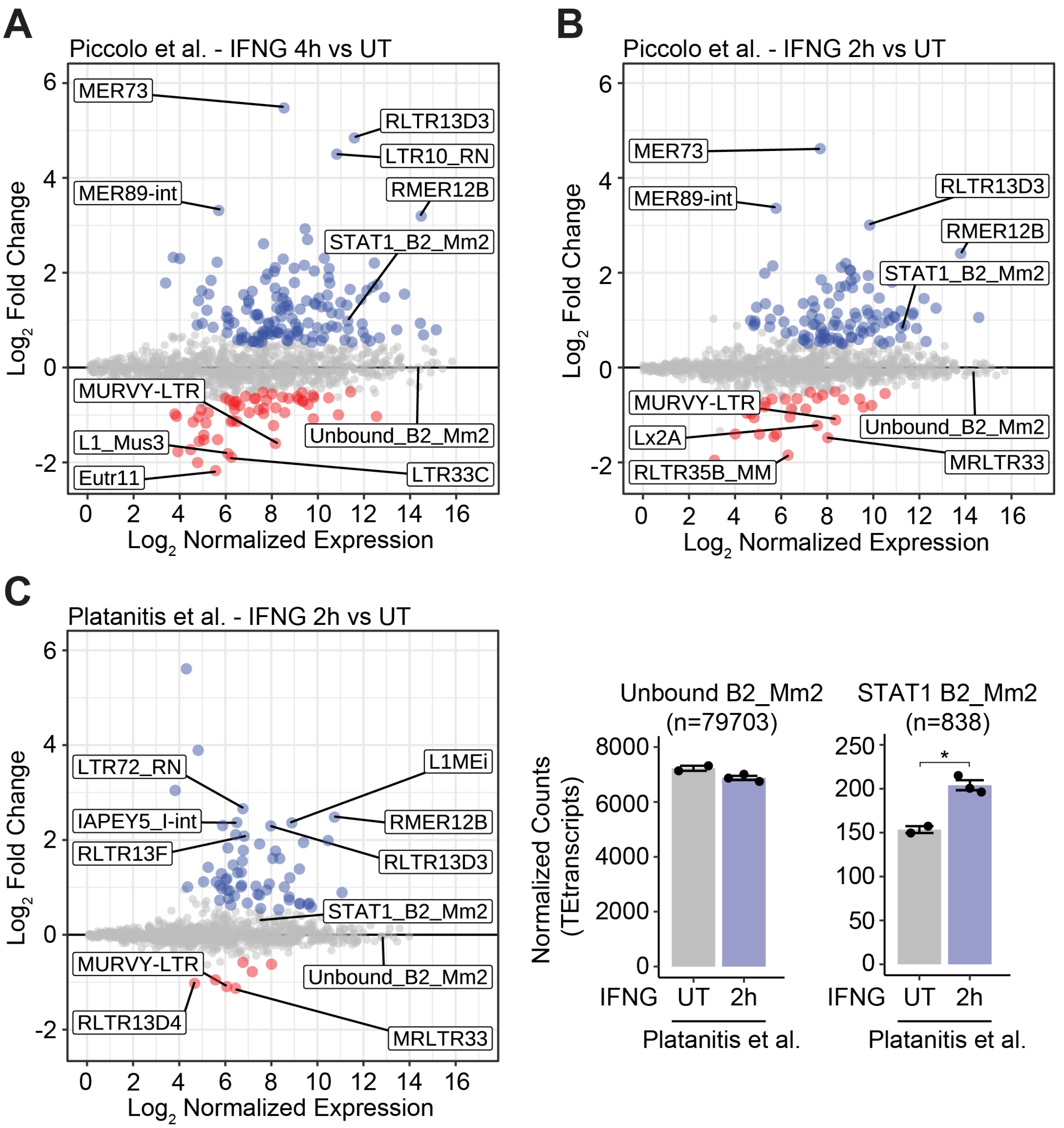


**Fig. S2.** IFNG-inducible TE expression in murine BMDMs. (A) MA plot visualizing immune-stimulated changes in TE family-level expression according to RNA-seq from murine BMDMs stimulated with IFNG for 4 hours. Unbound B2_Mm2 (n=78419) and STAT1-bound B2_Mm2 (n=2122, defined through Piccolo et al. (1) IFNG 2 hour ChIP-seq) are designated separately. Significantly downregulated (log2FC < -0.50, FDR < 0.05) genes are shown in red while significantly upregulated (log2FC > 0.50, FDR < 0.05) genes are shown in blue. (B) Same as in (A) but using RNA-seq from BMDMs stimulated with IFNG for 2 hours. (C) MA plot (left) and DESeq2 normalized counts (right) visualizing immune-stimulated changes in TE family-level expression according to RNA-seq from murine BMDMs stimulated with IFNG for 2 hours. Unbound B2_Mm2 (n=79703) and STAT1-bound B2_Mm2 (n=838, defined through Platanitis et al. (2) IFNG 1.5 hour ChIP-seq) are designated separately. Individual points are colored as in (A). Normalized counts shown for untreated (n=2) BMDMs and BMDMs stimulated with IFNG for 2 hours (n=3). **DESeq2 *p*-value < 0.001. Error bars designate SEM. BMDM: Bone marrow derived macrophage.

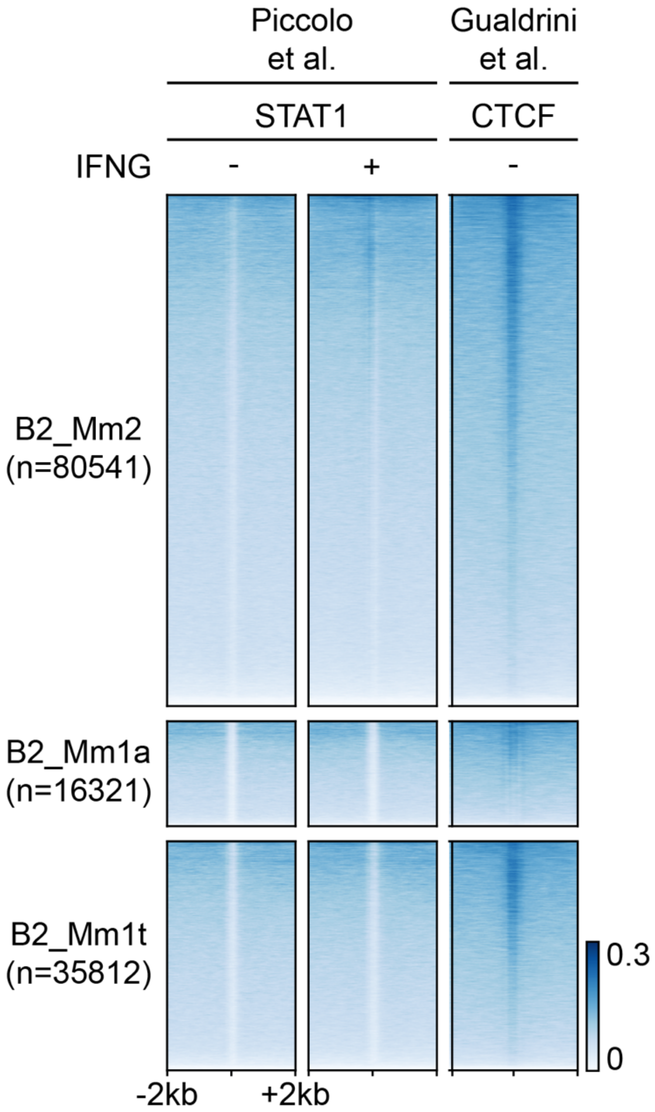


**Fig. S3.** STAT1 and CTCF occupancy overlapping all B2 elements in murine BMDMs. Heatmaps showing CPM-normalized ChIP-seq signal over B2_Mm2 (n=79073), B2_Mm1a (n=16223), and B2_Mm1t elements (n=35428) over a 4kb window. Elements without overlapping ChIP-seq signal were excluded. Regions are sorted by descending mean CPM signal. Signal intensity is indicated to the right. STAT1 tracks derived from Piccolo et al. (1). CTCF track derived from (3).

```
B2_Mm2.Dicer1     ----CTGGAGAGATGCCTTAGTGGTTAAGAGCA-CTGACTGCTCTTCCAGAGGTCCTGAG
B2_Mm2            GGGGCTGGAGAGATGGCTCAGCGGTTAAGAGCA-CTGACTGCTCTTCCAGAGGTCCTGAG
B2_Mm1a           GGGGCTGGTGAGATGGCTCAGTGGGTAAGAGCACCCGACTGCTCTTCCGAAGGTCCGGAG
B2_Mm1t           GGGGCTGGTGAGATGGCTCAGTGGGTAAGAGCACCCGACTGCTCTTCCGAAGGTCCGGAG
                      **** ****** ** ** ** ******** * ************  ****** ***

                       GAS      CTCF                                    CTCF
                       TTCCCAGAA CCACAAGGTGGC                           GCCCTC
B2_Mm2.Dicer1     TTCAATTCCTAGAAACCATGTGGTGGCTCACAATCATCTGTAATGGGATCCGATGCCCTC
B2_Mm2            TTCAATTCCCAGCAACCACATGGTGGCTCACAACCATCTGTAATGGGATCTGATGCCCTC
B2_Mm1a           TTCAAATCCCAGCAACCACATGGTGGCTCACAACCATCCGTAACGAGATCTGACTCCCTC
B2_Mm1t           TTCAAATCCCAGCAACCACATGGTGGCTCACAACCATCCGTAACGAGATCTGACGCCCTC
                  ***** *** ** *****  ************* **** **** * **** **  *****

                  CTCF
                  TTCTGG
B2_Mm2.Dicer1     TTCTGGCGTGTCTGAGGACAGTGACAGTGTACTCATATG----GTAAAATAAATACATCT
B2_Mm2            TTCTGGTGTGTCTGAAGACAGCTACAGTGTACTCACATACATAAATAAATAAATAAATCT
B2_Mm1a           TTCTGGAGTGTCTGAAGACAGCTACAGTGTACTTACATAT---AATAAATAAATAAATCT
B2_Mm1t           TTCTGGTGTGTCTGAAGACAGCTACAGTGTACTTACATAT---AATAAATAAATAAATCT
                  ****** ******** *****  ********** * **        ********* ****


B2_Mm2.Dicer1     T--------AAAGAAG
B2_Mm2            TTAAAAAAAAAAAAAA
B2_Mm1a           TTAAAAAAAAAAAAAA
B2_Mm1t           TTAAAAAAAAAAAAAA
                  *        *** **
```

**Fig. S4.** Multiple sequence alignment of consensus sequences for B2_Mm2, B2_Mm1a, and B2_Mm1t as well as the sequence for a B2_Mm2 element intronic to Dicer1. Predicted GAS and CTCF motifs (FIMO *p*-value < 1×10-4) are indicated above. Base changes relative to the motif consensus corresponding to poorly weighted positions in the motif position weight matrix are highlighted in green. Base changes in positions that are canonically important for transcription factor binding and weighted heavily in the position weight matrix are highlighted in red.

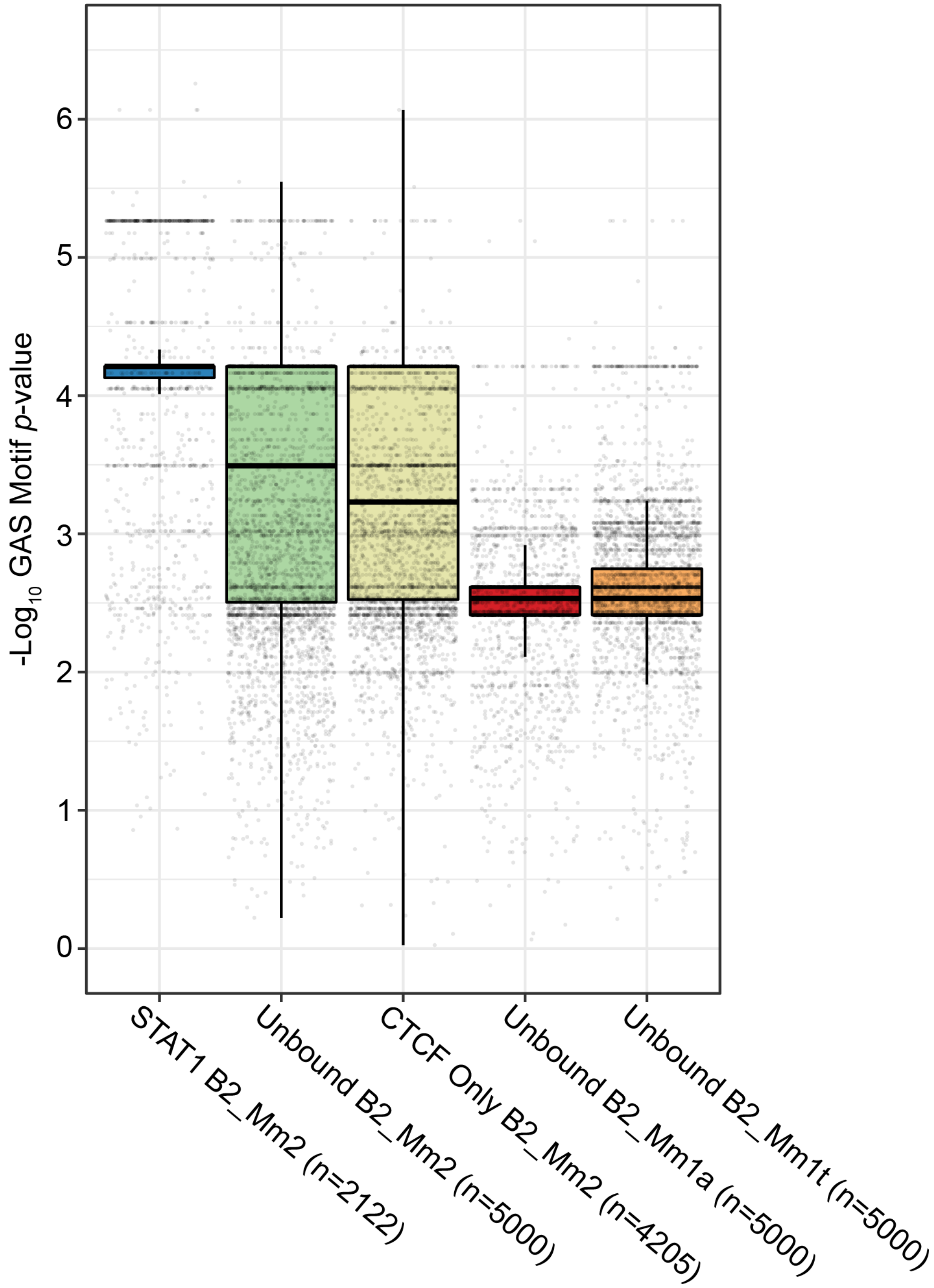


**Fig. S5.** Box-and-whisker plot visualizing the distribution of FIMO *p*-values for GAS motifs overlapping B2 elements. B2 elements were classified as follows: STAT1-bound B2_Mm2 (n=2122, blue); B2_Mm2 not bound by CTCF nor STAT1 (n=5000, green); B2_Mm2 bound by CTCF but not STAT1 (n=4205, yellow); unbound B2_Mm1a (n=5000, red); and unbound B2_Mm1t (n=5000, orange). Each dot represents the *p*-value for the most significant predicted GAS motif overlapping one B2 element. The median, lower quartile, and upper quartile are as indicated.

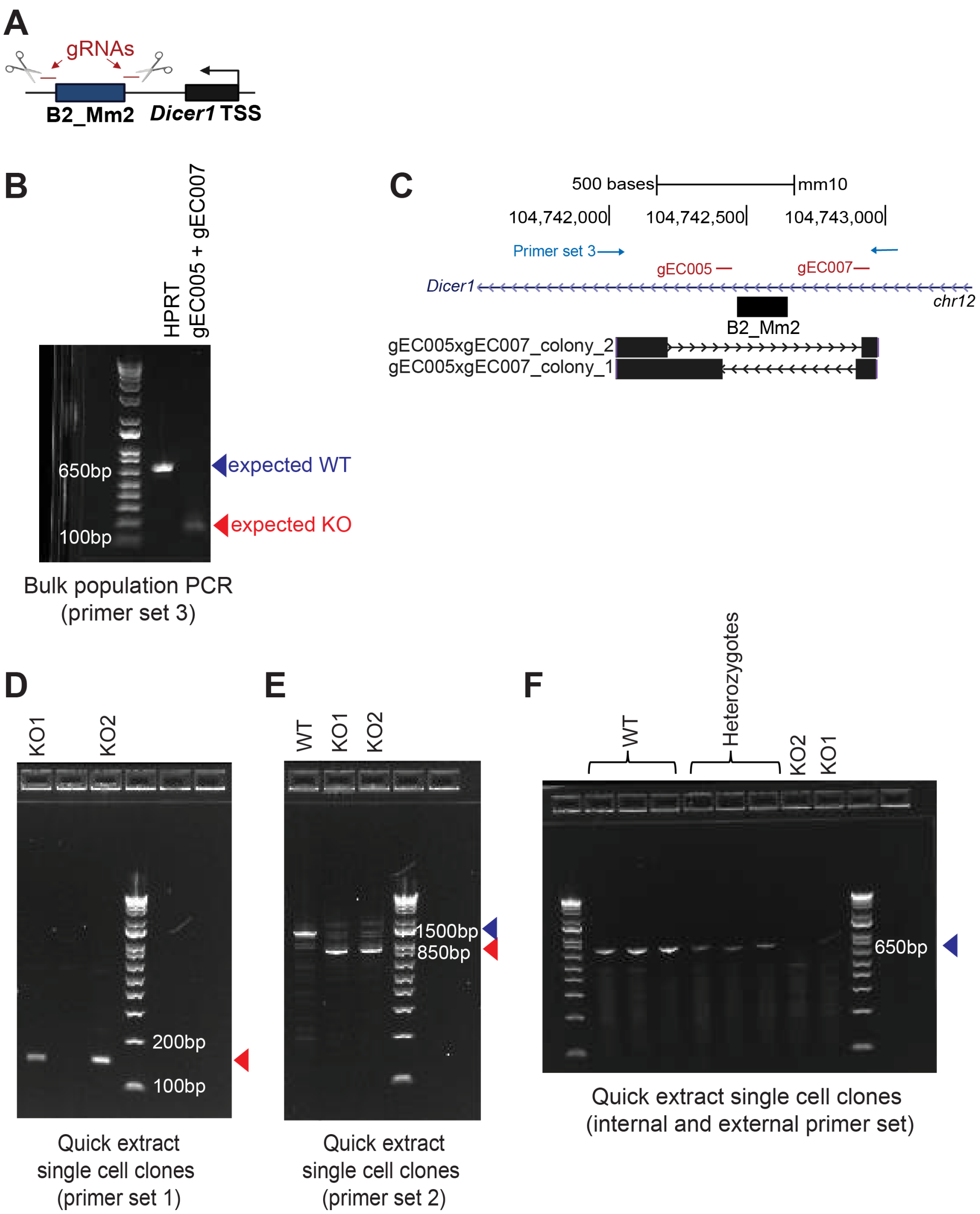


**Fig. S6.** Validation of CRISPR B2_Mm2 knockout in J774 cells. (A) Schematic showing design of guide RNAs (gRNAs) flanking the B2_Mm2.*Dicer1* element. (B) PCR validation of CRISPR/Cas9-mediated knockout of B2_Mm2.*Dicer1* in bulk J774A.1 cells. HPRT- serves as a control with an expected WT band at 627bp. Successful KO is shown as indicated by the expected gEC005xgEC007 amplicon at 127bp. (C) Genome browser track showing Sanger sequencing results from the bulk cell population aligned using BLAT (4). Sequencing of different E. coli clones derived from the same bulk cell population reflects presence of two distinct KO alleles. Positions of gRNAs and PCR primers are indicated in red and blue, respectively. (D, E) PCR validation of B2_Mm2.*Dicer1* KO in clonal J774A.1 cells using two distinct primer sets external to B2_Mm2.*Dicer1*. (F) Same as in (D) and (E) but using a pair of internal and external primers.

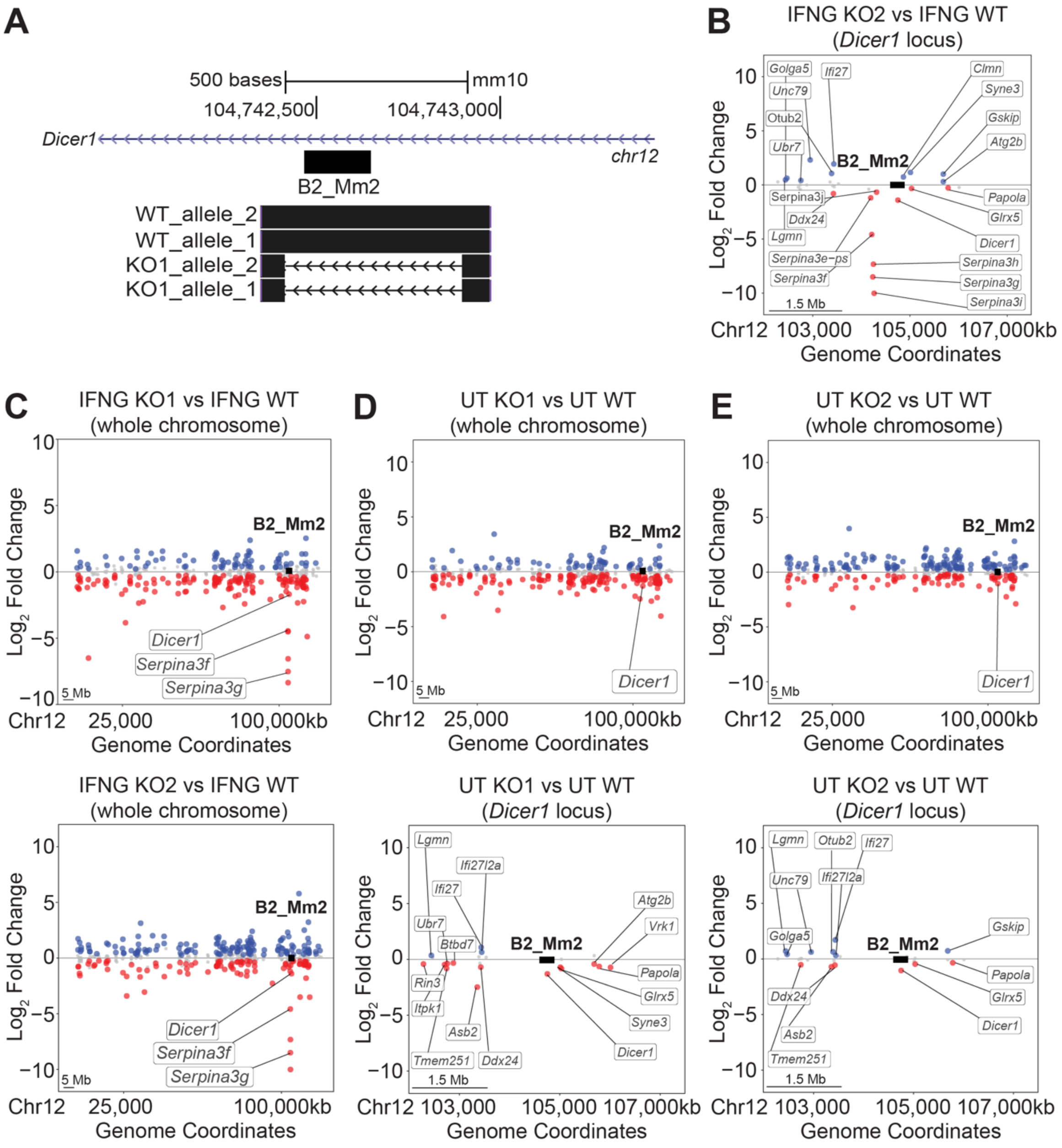


**Fig. S7.** B2_Mm2.*Dicer1* in the genomic landscape. (A) Genome browser track showing Sanger sequencing results from WT and KO1 J774A.1 cells aligned using BLAT (4). (B) Distance plot showing changes in gene expression within a 5Mb window of B2_Mm2.*Dicer1* in IFNG KO2 cells relative to IFNG WT. (C) Distance plot showing changes in gene expression across Chr12 in IFNG KO1 cells relative to IFNG WT (top) and in IFNG KO2 cells relative to IFNG WT (bottom). (D) Distance plot showing changes in gene expression UT KO1 cells relative to UT WT across Chr12 (top) and within a 5Mb window around B2_Mm2.*Dicer1* (bottom). (E) Distance plot showing changes in gene expression UT KO2 cells relative to UT WT across Chr12 (top) and within a 5Mb window around B2_Mm2.*Dicer1* (bottom). (B-E) Significantly downregulated (log2FC < 0, FDR < 0.05) genes are shown in red, and significantly upregulated (log2FC > 0, FDR < 0.05) genes are shown in blue. B2_Mm2.*Dicer1* is represented as a black box (not drawn to scale). UT: Untreated. KO: Knockout.

**Table S1 (separate file).** DESeq2 & gene ontology results for murine BMDMs genes stimulated with IFNG for 2 or 4 hours.

**Table S2 (separate file).** Murine BMDM ChIP-seq motif enrichment using XSTREME.

**Table S3 (separate file).** Coordinates for STAT1-bound TEs in murine BMDMs.

**Table S4 (separate file).** Family-level TE enrichment over murine BMDM ChIP-seq using GIGGLE.

**Table S5 (separate file).** DESeq2 results for TE expression in murine BMDMs stimulated with IFNG for 2 or 4 hours.

**Table S6 (separate file).** Differential B2_Mm2 motif enrichment analysis using AME.

**Table S7 (separate file).** Absolute distances for STAT1-bound B2 elements to the nearest BMDM ISG.

**Table S8 (separate file).** Excel spreadsheet of all standard primer, qPCR primer, and gRNA sequences.
